# Measuring researcher independence using bibliometric data: A proposal for a new performance indicator

**DOI:** 10.1101/388678

**Authors:** Peter van den Besselaar, Ulf Sandström

## Abstract

Bibliometric indicators are increasingly used at the individual level – as is exemplified by the popularity of the H-index and many other publication and citation based indicators used in evaluation. The issue isn’t whether these indicators can be considered useful, as they do provide a description of a researcher’s oeuvre. However, at the same time, they are not enough to assess the quality of a researcher and his/her oeuvre: Quality has more dimensions than productivity and impact alone. In this paper, we argue that independence is an equally important characteristic that however lacks validated indicators for measuring it at the individual level. We propose two indicators to measure different dimensions of independence: one assessing whether a researcher has developed an own collaboration network, and another assessing the level of thematic independence. We illustrate how these indicators distinguish between researchers that are equally productive and have similar impact. The independence indicator is a step forward in evaluating individual scholarly quality: in cases where citations and publications do not distinguish, the indicators for independence may do.

## Introduction

The use of (bibliometric) indicators for evaluating research performance has become rather common, at the level of research organizations and research teams, but after the development of the H-index also at the individual level. Over the years, the number of indicators is proliferating, and a recent paper even distinguished 108 indicators at the individual level [1]. However, despite the large number, almost all of these indicators are focusing on productivity (publications) and impact (received citations), and therefore measure in many different ways productivity and scholarly impact, as is visible in recent work on individual level metrics [2, 3, 4, 5]. Only a few of the 108 indicators are focusing on another quality dimension, namely collaboration. Other reviews of individual level indicators support this. In a study on the relations between bibliometric indicators, Bollen and colleagues did a principal component analysis of 38 citation and usage indicators [6]. The types of indicators included are citation based, but also indicators derived from download data (click stream data). The principal component analysis indicated two dimensions of impact: (i) short term impact (usage) versus long-term impact (citations), and (ii) popularity versus prestige. Overall, many indicators focus on scientific impact of papers and related on the impact of journals, and only a few network-type indicators have been developed, using the co-author network to measure an author’s impact [1, 7].

In parallel, a discussion has emerged amongst evaluators and bibliometricians about the reliability and validity of the indicators, and increasingly criticism is formulated of the indicators and their use [8, 9, 10]. The critique can be summarized as follows: (i) peer review should remain the main procedure, and bibliometric indicators can at best inform peer review^1^; (ii) the quality of indicators needs permanent scrutiny; (iii) evaluation should relate to the goals; and (iv) indicators have perverse effects. More specifically, as bibliometric data have become available widely, ‘everybody’ is able to calculate indicators, which is seen as very risky, as most non-professionals do not understand the meaning of the indicators, and apart from that, are not able to calculate the indicators in a correct way, e.g. to field-normalized citation impact [11].

Most attention has been on the so-called perverse effects^2^, but here we focus on the other main point, the validity of the indicators: Do bibliometric indicators measure what we want to measure? As already discussed, most indicators focus on high productivity, high impact, and a high-quality collaboration network.^3^ If these are of restricted value for a more inclusive measurement of scholarly quality, we may need additional ones. The starting point can be to systematize the various quality dimensions that should be taken into account when hiring academic staff, or when selecting research proposals for funding. In order to enrich the set of useful indicators, it would be good to study in more detail what constitutes a high-quality researcher (for an interesting account, see [12]).

The discussion on metrics has in fact been rather restricted and misses the link to the larger evaluation frameworks that are or might be used, on the institutional and group level, but also for assessing researchers for hiring and tenure and promotion [13]. For a previous study we interviewed some 40 panel members, who mentioned a large and diverse set of different quality dimensions they used in evaluation [14]. The panel members made a distinction between criteria for evaluation grant applications, and for jobs. Several dimensions could be distinguished: (i) Productivity proved to be a main criterion, as earlier grants, originality, a broad scope, and independence; (ii) other work oriented personal characteristics were mentioned too – like commitment and hard-working, and ambition, and when it is about academic jobs, also (iii) social characteristics – like being a pleasant person, and a team player (Table 1).

**Table 1:**
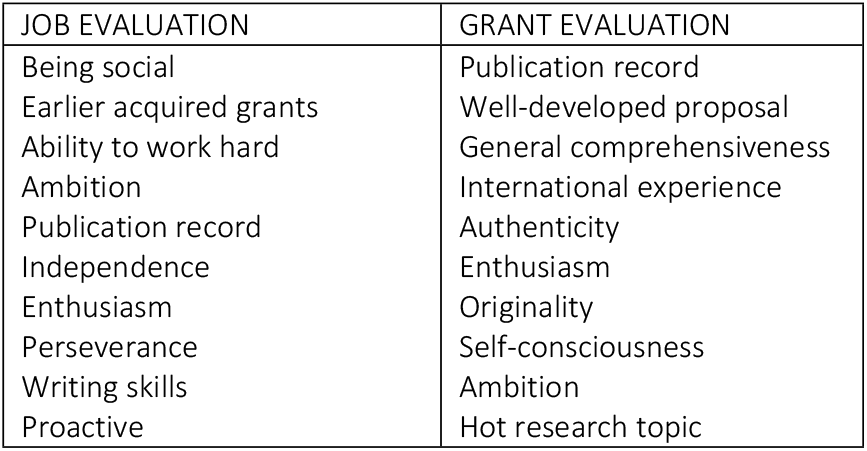
Evaluation dimensions

Interestingly, the tendency in many debates seems to be that less use of indicators is the better alternative for low valid indicators. Here we follow a different strategy: if current indicators are not covering the various quality dimensions and lack validity, let’s develop better indicators for those dimensions of quality not yet addressed. This seems more relevant than trying to marginally improve the existing one’s [15].

One of the criteria mentioned for jobs was independence. As science develops through original and new ideas, there is a need for independent researchers that challenge the existing common views and open up new lines of research. Independence, in this sense, is breaking with existing lines of work and coming up with something new. To achieve higher academic positions, and to get prestigious grants, one should have shown this type of independence. The question of independence gets an additional relevance through the growth of research collaboration and team science [16, 17, 18]. If most work is collaborative, this must have implications for the assessment processes and the indicators deployed. An independence indicator may be one of the instruments to disentangle the performance from complex network dependencies. It helps to fill the gap in the existing set of quantitative indicators [13].

## Independence

That independence is important can be illustrated in various ways. A report published by the American Academies of Science [19, see also: 20] actually points out that the conditions for becoming independent were deteriorating: the age at which researchers got the first independent grant in the US was increasing. This would imply that researchers are becoming independent too late, something that also may work negatively on attracting new generations to scientific research. Although this is a different topic than the one we discuss in this paper - how to recognize independence - it does show that independence of researchers is a relevant research policy issue. Because requests for grants from new generation investigators were evaluated on the basis of “preliminary results”, most funded research became constrained to well-worn research paths — those previously pursued by the new investigators when they were postdoctoral fellows in established laboratories. In short, innovation was the victim of a system that had become much too risk adverse:

> “Furthermore, the study sections want significant assurance that the project will work, and so they require the principal investigator to have surmounted at least some of the experimental hurdles inherent in the project prior to funding. Reasonable young principal investigators are quick to get the message: stay within the confines of known systems and proven technologies, and do not challenge existing beliefs and practices.” [20]

Consequently, there is a serious concern that new investigators are being driven to pursue more conservative research projects instead of the high-risk, high-reward research that more likely advance science significantly. The special creativity that younger scientists may bring in may also get lost when these investigators are forced to focus on others’ research. In this way, we define an “independent investigator” as one who enjoys independence of thought—the freedom to define the problem of interest and/or to choose or develop the best strategies and approaches to address that problem. Under this definition, an independent scientist may work alone, as the intellectual leader of a research group, or as a member of a consortium of investigators each contributing distinct expertise. “Independence” does not mean necessarily “isolated” or “solitary”, or imply “self-sustaining” or “separately funded” [19, p21].

A good example is the Starting Grants scheme of the ERC [21, 22]. The ERC instructs its panels to select excellent researchers and groundbreaking projects, and one of the few concrete criteria they mention is ‘independence’, defined as having one (or a few) publications without the former PhD supervisor being co-author.^4^ When interviewing the ERC panelists, independence indeed proved a major issue, but how it can be measured remained unclear. Some would argue that single authored articles indicate independence^5^, but in a world where one hardly can do research alone, this becomes a slightly irrelevant indicator.

That bibliometric indicators have limitations in this respect has been pointed out by many [23, 24]. It was also the focus for a working party of the Swedish medical research council. Their final document states:

> “The assessment of independence is a frequently discussed issue, especially when it comes to researchers in large collaborative projects and/or young researchers. (…) Given that research today often includes networks and large collaborative research teams, it may be difficult to discern the independence of individual researchers and especially of the young. A young researcher in a network often shows many publications, but without a prominent place in the author list, which complicates the assessment of the degree of independence.” [26, our translation]

The importance of independence in the Swedish setting is also illustrated by the longstanding work of docent promotion committees at faculties of medicine. For several years a working group of these committees has been trying to operationalize the rules. One example how they treat independence is the Karolinska Institute’s regulations for those committees:

> “Independence can be documented, for example, when the applicant has been obviously senior representative of recent contributions to research, the latter not seldom exemplified by the recent scientific publications, and has conducted his [sic!] own consistent research line after the dissertation. Independence can also be shown through research responsibilities as a supervisor for a PhD student or a guest researcher.” (Instructions for *docent position*^6^ at Karolinska Institute, established by the Vice-chancellor March 18, 2014)

If independence is an important evaluation dimension, the question is whether we can develop useful indicators to measure independence in a reliable and valid way. This is the aim of this paper.

Despite the fact that many would consider independence as a main criterion for selection, it is not easy to use. For example, when female researchers are evaluated they are often considered as “dependent” on their supervisor while male applicants more often were considered as “independent”, even when they followed in the research lines of their supervisor [26]. A recent qualitative study performed in Sweden found the same pattern [25]. Results were based on direct observation of committees inside the Swedish Research Council.

Findings related to “Independence” are reported in a special section of the committee evaluation report. The report issued by NAS [19] suggests the need for an indicator that could be used to measure independence as a more valid measure, reflecting the meaning of the concept and getting it uncoupled from the many gender biased or other stereotypes:

First of all, a successful researcher of course needs to have acquired excellent research skills and produced relevant results, which can be measured using publications and citations. However, as most research is teamwork, these publications and citations could have been ‘borrowed’ from an excellent team, in which a researcher not necessarily has had the leading role. Therefore, to get tenured or to get a prestigious grant, an early career researcher should also have developed independence. To be successful as a researcher, one needs to be able to formulate one of preferably several own, independent and promising research lines. This would result in papers on topics that are different from those of the environment where the early career was spent, and with other co-authors than supervisors and colleagues from that period. The aim of this paper is to develop an operational concept of independence, and to provide a proof of concept. We will define the indicators and show how ‘independence’ can be measured using data about the co-author network and the publications network.

Independence is not the only relevant dimension. One may also think of ‘originality’, ‘risk taking’, or ‘interdisciplinary’ as important features of the research lines to be developed [22, 27]. And the own research line should be promising and relevant, so probably within a research field that shows growth. Whether the selected own research lines are promising should also be measured: with traditional indicators like top cited papers, but also using science maps based indicators that position the research lines within the larger landscape: is this a line within a hot or cold field? Is attention for the field growing and if so, fast? Sandström & Sandström [2] and Wang [28] are but two examples of methods for finding hot areas that are based on similarities between papers. This would lead to several other relevant new indicators, which we will not develop here, but briefly mention them as part of our wish list for better indicators [27, 29, 30]. Here we focus on our core independence indicators.

## Independence indicators

An independent researcher, in the line of the discussion above, has taken the freedom to define the problem(s) he/she want to address, and to choose the best strategies and approaches to address that problem(s). Independence in this definition is something with a time dimension: independence emerges over the years. It implies in the first place that a young researcher becomes independent from the environment where he/she was educated. As PhD student, and as Postdoc, a researcher works in projects designed by others, often the PI who also functions as supervisor. After having done that for a while, the researcher should have become skilled and experienced to formulate his/her own research questions and projects at challenging research fronts, unguided by the former supervisors. And he/she should try to get resources to do so. This, of course, does not mean that independent researchers avoid collaboration, as almost all research nowadays is collaborative. In fact, building up relevant collaboration networks is one of the resources needed for excellent research. The challenge is to formulate indicators that can tease out the level of independence from a network of dependencies^7^.

### Indicator 1: The structure of the co-author network

If independence means that a researcher does not depend anymore on the environment where he/she was trained as researcher, we would expect to see that in the collaboration network. An independent researcher is expected to collaborate with a variety of others, but less so with the former supervisor(s). Therefore, independence can be defined as a quality of the co-author network of the researcher. The size of the network indicates how the environment of a researcher perceives his or her contribution. The more someone has to contribute, the more other researchers want to collaborate, so the more co-authors someone has. A young researcher is often introduced in the academic world through the supervisor. In the beginning of the career, the co-author network of a young researcher is therefore expected to be embedded in the network of the supervisor; something that may be helpful in the first career steps. But after a while, the co-author network of an independent researcher will significantly differ from the supervisors’ co-author network. This can be measured using two network properties of the co-author *ego-network* of the researcher: (i) the *eigenvector centrality* of the former supervisor in the co-author ego-network of the researcher, and (ii) the *clustering coefficient* of the former supervisor in the ego-network of the researcher.

The *eigenvector centrality* expresses influence in the network: in contrast to the degree centrality, the eigenvector centrality takes into account the connectedness of other nodes. A node is important (the eigenvector centrality is higher) if it is connected to other important nodes. So even if the supervisor is not connected to all nodes in the researcher’s network, if he/she is connected to the important (highly connected) nodes, the supervisor’s eigenvector centrality is still high. The *clustering coefficient* measures the extent to which a node is part of a clique in the larger network. The more a supervisor is part of a specific clique within the researcher’s network, the less the researcher’s network coincides with the network of the supervisor, suggesting higher independence.

The researcher is the center of his/her own ego-network, and will therefore have a eigenvector centrality of 1. The more independent a researcher is, the more distinct coauthor cliques he/she will be part of, and that is expressed in a low (down to 0) clustering coefficient. If the two network scores of the supervisor are similar to those of the researcher, independence is low; the more the researcher’s network is his/her own, the lower the eigenvector centrality (approaching 0) and the higher the clustering coefficient (approaching 1) of the former supervisor will be.

The proposed indicator may not work in all research fields. If there is only one co-author clique (clustering coefficient = 1), it would be impossible to assess the individual independent contribution of the researcher under evaluation. This may be the case in research fronts where ‘everyone’ is on all papers (hyper-authors) as in some physics subfields. Also in fields where single-authored papers are the dominant mode, the indicator would not work, as co-author networks do not exist. However, in all fields, the share of coauthored papers grows fast [31].

### Indicator 2: The cognitive network of the researcher

Being independent also means that the researcher starts to explore new topics and to develop own research lines, by following his or her own ideas. The researcher should move to new research questions not belonging to the research agenda of the former supervisor. This explains why the number of citations and publications may not be decisive in assessing the performance of researchers. The real issue is whether one publishes and is cited because of one’s own good research, and not because of the good performance history of the supervisor. This implies that after a while, the publications of the early career researcher should be outside the research front(s) that the former supervisor’s publications belong to.

To measure whether the researcher developed own research lines, independent of the former supervisor, we downloaded from the Web of Science (Online version) the papers of the researcher and the former supervisor, including their co-authored publications. We select all papers published until two years after the main career decision: appointment as tenured (associate) professor versus leave academia. The additional two years are included to account for papers that were written and possibly even accepted, but not yet published before promotion/leaving dates.

We created the joint paper network of researcher and supervisor using bibliographic coupling. This results in a network of several components and clusters representing different strands of research. Is an own research line of the researcher visible in the network? Or are the own papers of the researcher included or even ‘hidden’ in the network of the supervisors’ papers? If the latter is the case, the researcher has remained within the research program of the supervisor, and no own program was developed. For the joint network, we calculate a similarity indicator based on bibliographic coupling between the (partly overlapping sets of) papers of the supervisor(s) and of the researcher. The similarity measure is based on Salton’s Cosine Index and varies between 0 and 1 [32]. In order to account for research lines, we use only the following document types: Articles, Letters, Proceeding Papers and Notes. Reviews are, in our understanding, not representations of an individual researcher’s research line as there are many references to research that might be remote to the researcher in question. The SML algorithm (with resolution set to 1.5) was used to detect and break down the set of papers into topical clusters [33].

Some clusters contain only papers of the supervisor, others only papers of the researcher and again other clusters contain papers of both, as well as co-authored papers. If a researcher developed his own research line, which would imply that the researcher explores other/new questions, and refers to different literatures, then the similarity will be lower; if he/she continues within the research line of his supervisor, the similarity measure will be higher.

We calculate this independence indicator in the following way: we divide the number of all topical clusters where the researcher is active in but the supervisor(s) is not^8^ by the sum of these own clusters and the joint clusters of the researcher and the supervisor(s). The indicator value is between 0 (if the researcher has only been active in topics in which the supervisor also has been active at some moment) and 1 (if the supervisor is not visible in any of the researchers’ topical clusters).

## Data

This paper intends to be *a proof of concept*, and we apply the indicators on four researchers and their supervisors to show how it works. Measuring independence using these indicators can be done for any researcher given that the person in question has publications covered in citation indexes, e.g. Web of Science. However, we are interested in the question whether the indicator is valid, that is whether it measures independence as taken into account by e.g., panels that select applications for positions or jobs. In order to do that one needs pairs of researchers which are similar in various dimensions (age, field, talent, gender, academic performance in terms of publications and citations, etc.) but differ in independence. Having a few of such pairs, we can give a tentative answer on the question whether the measured independence relates to e.g., career success.

In another study [34] data of this type were collected in order to compare relatively homogeneous pairs of researchers. Research managers of several universities were asked for pairs of researchers that were seen as equally very talented and promising in the beginning of their careers but of which only one of the two had a successful academic career whereas the other left the academy. In this way pairs were created where both researchers were very talented and in the same subfield. For the current study, we used two of these pairs. Researchers A and B are in the same chemistry field; researchers C and D are in the life sciences. Researchers A and C became full professor, and researchers B and D left the university for a position in industry. For all researchers, the decisive career moment was in their late thirties, some nine years after receiving their PhD degree.

Back to our current paper; we downloaded the bibliometric data (WoS) of the publications of the researchers, as well as those of their PhD supervisors, and checked manually for false positives and false negatives. The WoS data cover the career from the start of the PhD research up to their late thirties. For the supervisor(s) we also include publication data of a decade before the start of the PhD trajectory of the researcher, in order to cover the relevant research portfolio. As Fig 1 shows, all four researchers had quite a good performance level.^9^ If publications and citations would be decisive, one would expect RB to have a better academic career than RA, as RB had a much higher impact. The same holds for RD compared to RC.

**Fig 1:**
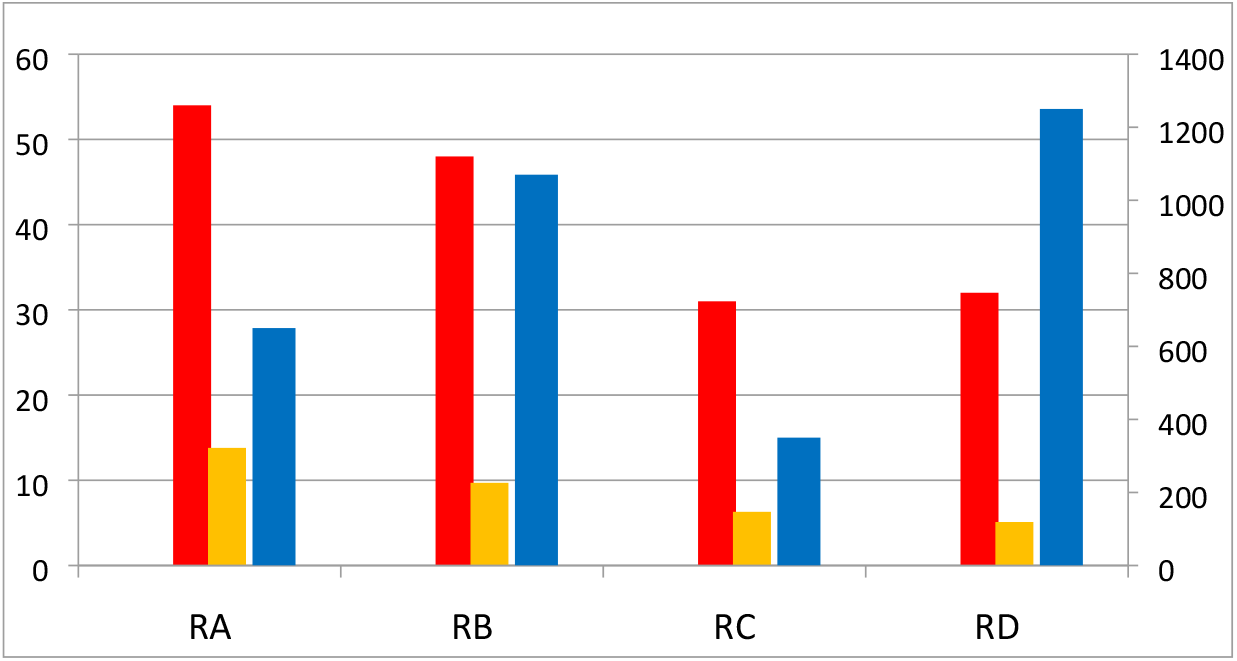
Performance scores of the five researchers at decisive career moment. Publications: red (full), orange (fractional) = left axis; citations: blue = right axis. Source: WoS

The following analyses were done for the period between the PhD period and the moment of the main career decision:

1. Calculation of the share of papers *co-authored* with the supervisor. The lower the score, the less support the researcher may have had from the supervisor. The higher the score, the less autonomous the researcher may have become.
2. Calculation of *research line similarity* between researcher and former supervisor(s) over a period from the start of PhD until the main career decision.
3. Calculation of the eigenvector centrality and the clustering coefficient for the supervisors of the researchers in the *ego network* of the researcher.
4. Then these calculations are *combined* into an overall independence indicator.

## Findings

### Pair 1

We first compare RA and RB, both in chemistry. One of the researchers had a successful academic career (RA with supervisor SA) and became full professor. The other researcher (RB with supervisor SB) left the university for an industrial R&D lab – so productivity and impact were not decisive here (Fig 1). Table 2 shows the performance of the two researchers at three moments: the moment of the PhD, the moment of the decisive career decision, and ten years later. In terms of performance, RB had more publications and more citations than RA. Interestingly, about ten years after RB left academia, the H-indexes of both are equal, showing that RB’s work is still appreciated by the community.

**Table 2:**
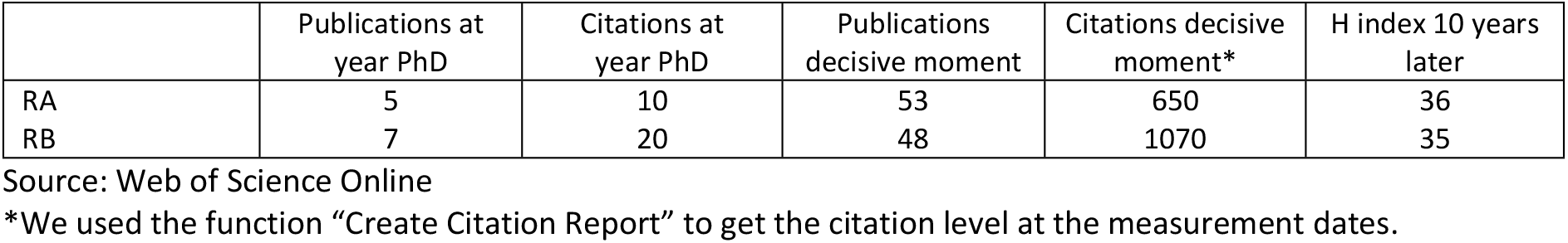
Performance of RA and RD, three moments in time

|  | Publications at year PhD | Citations at year PhD | Publications decisive moment | Citations decisive moment* | H index 10 years later |
| --- | --- | --- | --- | --- | --- |
| RA | 5 | 10 | 53 | 650 | 36 |
| RB | 7 | 20 | 48 | 1070 | 35 |

Source: Web of Science Online
\*We used the function "Create Citation Report" to get the citation level at the measurement dates.

In the relevant period, RA and RB have about the same number of co-authors (5.1 versus 5.2, respectively). Both researchers co-authored with their supervisor. In case of RA, about 20 % of his publications are co-authored with his two supervisors, whereas RB co-authored about 95 % his papers with his supervisor SB, both in the period under consideration, so RB collaborated much more intensively with his supervisor SB than RA did with SA. We deliberately took a pair with strong co-authoring differences to illustrate the approach.

These different collaboration roles result in rather different network scores: the two researchers have in their own ego network by definition an eigenvector centrality of 1. Within the ego-network of RB, SB has a high eigenvector centrality (0.91), indicating that SB is almost as central in the network of RB as RB herself. In contrast, the eigenvector centrality of SA in the ego-network of RA is very low (0.09), indicating SA’s relatively marginal position in the network. Similarly, the clustering coefficient of SB is low (0.11), as low as RB’s clustering coefficient (0.09), but the clustering coefficient of SA is high (0.79), very different from the clustering coefficient of RA (0.07). This indicates that SA is connected to a specific subset of nodes only. Consequently, we may conclude that RB hardly has an own network, whereas RA does have one.

The other indicator measures the difference in research lines between the researcher and the former supervisor. Research lines are analyzed by creating a network of papers, based on bibliographic coupling. Papers cluster if they refer to a similar literature. One may calculate the similarity between two oeuvres as the average of the (cosine based) similarity between papers from the oeuvres – as explained in the methods section. Bibliographic coupling shows that the research lines of RA and SA differ, whereas the research lines of RB and SB are very similar. Visualization of the paper networks illustrates these findings.

Fig 2 shows the topics network of the researcher RA and the supervisors. We number the papers by topical cluster, so one may see the variety of topics worked on. The color of the nodes reflects authorship. The light blue nodes are supervisor 1, the dark blue supervisor 2, and the purple nodes are co-authored by the supervisors. The yellow nodes are authored by the researcher, and the green by the researchers plus the supervisor(s). Of course, in all cases, other co-authors may be involved.

**Fig 2:**
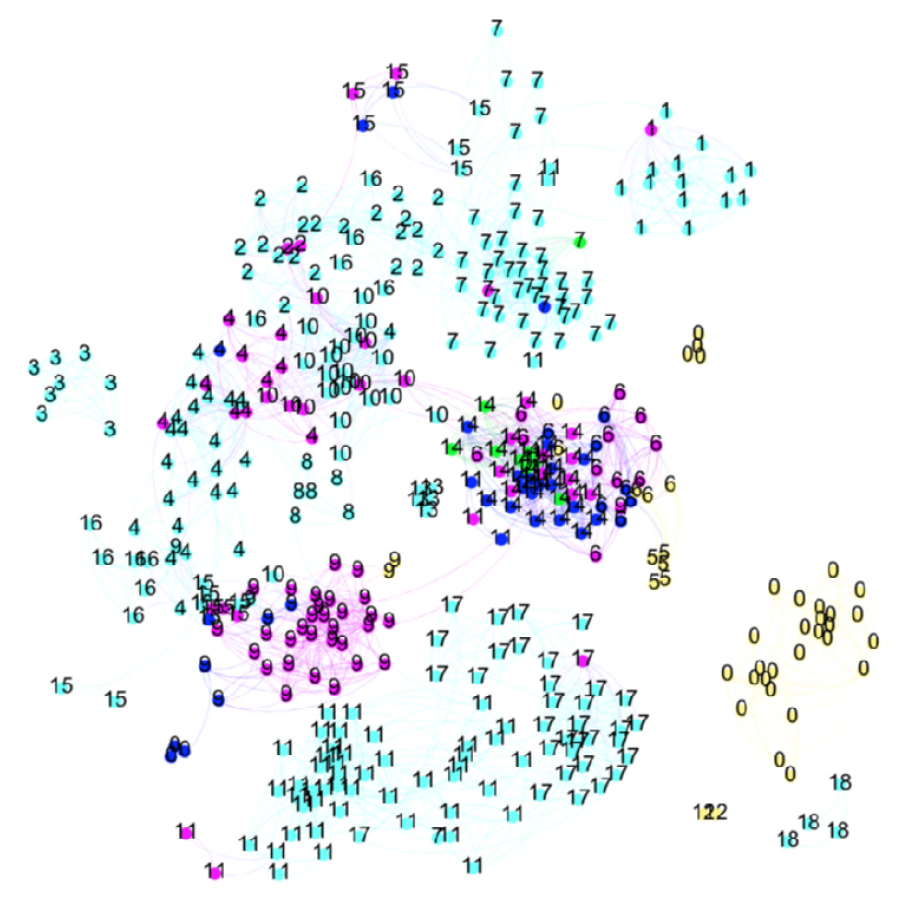
The topic network of researcher RA and two supervisors. Blue/purple = supervisor(s); Green = researcher co-authored with supervisor(s); Yellow: researcher Numbers indicate the topical clusters

Researcher RA worked intensively with supervisor 2 in topic T14 (middle of the map). Other joint clusters but with much less papers of the researcher are T6 and T9. But after receiving the PhD degree, RA started to work on other topics, in which neither of the supervisors has been active: T0, T5 and T12. Researcher RA has 83 % of the papers and 60 % of his research topics without participation of the supervisors. To investigate the further development of RA, we also created a bibliographic coupling map of the papers of RA and SA for another ten-year period (not included). RA remains detached from his former supervisor(s), and his own research lines have clearly developed extended over that period, unconnected to the work of former supervisor.

Fig 3 shows the similar information for RB and SB. Much more joint papers (green) are visible in Fig 3, and only two papers without the supervisor as co-author. These papers also form the only own topic of researcher RB (yellow nodes). All other papers of RB are strongly connected to clusters dominated by the former supervisor, and this has not changed over time, indicating the strong enduring similarity between their respective works.

**Fig 3:**
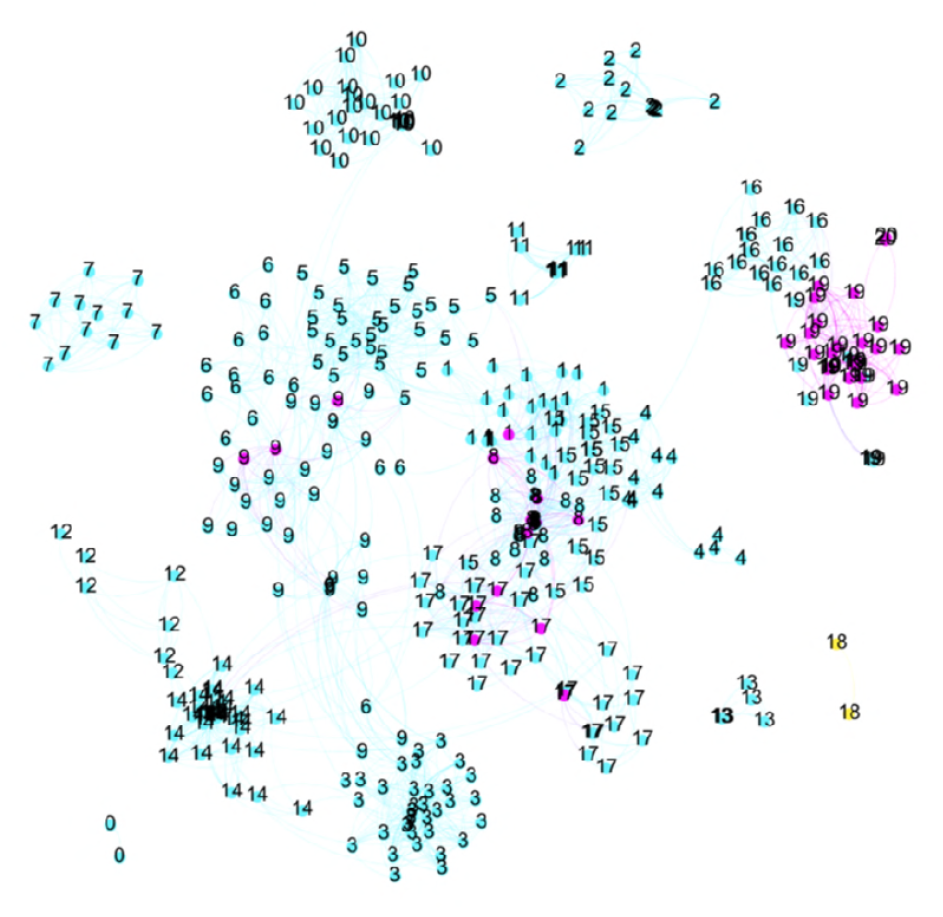
The topic network of researcher RB and supervisor

So, unlike RA, RB has hardly any work outside of the large network with SB: only 4 % of the papers and 14 % of the topics are without SB. RB, although productive and highly cited, did not develop new research lines but remained close to the work done with supervisor SB. Concluding, two similarly talented researchers in the same field and with about the same number of co-authors, productivity and impact, show strongly different patterns in their relation to the supervisor, and this is reflected in the scores for the proposed indicators. Fig 4 summarizes the findings.

**Fig 4:**
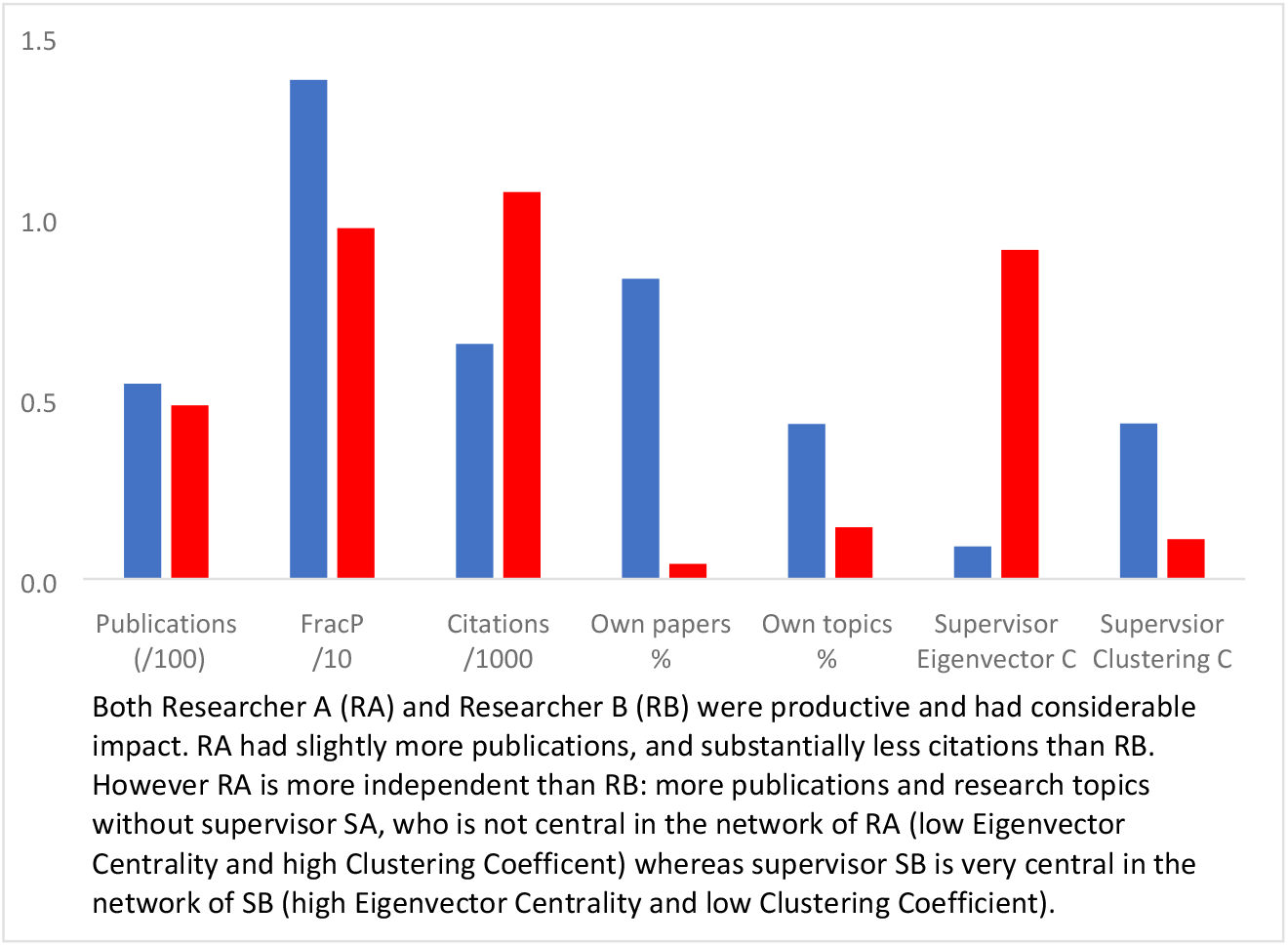
Indicator values for RA (blue) and RB (red)

### Pair 2

We now give the results for the second (life sciences) pair. As we showed in Fig 1, both researchers RC and RD were rather productive. RD had a higher citation impact. Also here they very differently co-authored with the supervisors. Researcher RD has much more intensively co-authored with the two supervisors (75% of the papers) than researcher RC (17%). The eigenvector centrality of SC in the ego-network of RC is very low (0.04) and the clustering-coefficient of SC is very high (0.76), showing that for researcher RC the role of the supervisor in the research network is very modest. For Researcher D, the pattern is more or less the opposite. Supervisor SD has a moderately high eigenvector centrality (0.28), showing that supervisor SD is more central in the ego-network of former PhD-student RD. The clustering coefficient of SD is also fairly high (0.36), and that seems to contradict with the high eigenvector centrality. However, both indicators are influenced by the specific collaboration pattern of RD who had two supervisors, but the two never co-authored a paper between them. The latter lowers the eigenvector centrality and increases the clustering coefficient of the main supervisor. In this case one may ‘merge’ the two supervisors, to get a better indicator for the position of the supervisors in the network of RD.

Is this collaboration pattern reflected in the research lines of the two researchers? As the different numbers of co-authored papers suggest, also the number of *own topics* of the researcher may differ. This is indeed the case: A large share of RC’s papers (0.83%) are not co-authored with the supervisor SC, and consequently half of the topics (50%) RC is active in are not covered by the supervisor. RD on the other hand has no research topics without one of the two supervisors (Fig 5).

**Fig 5:**
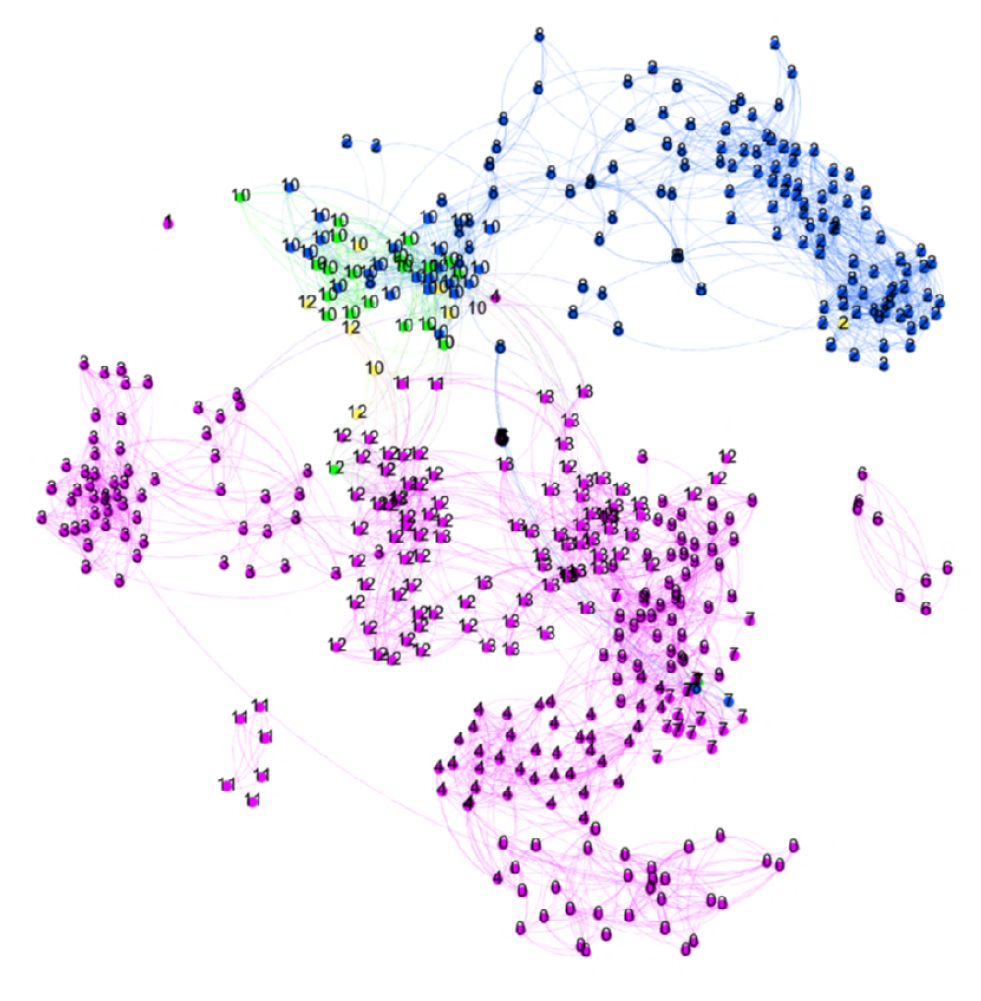
The topic network of researcher RC and the supervisors

**Fig 6:**
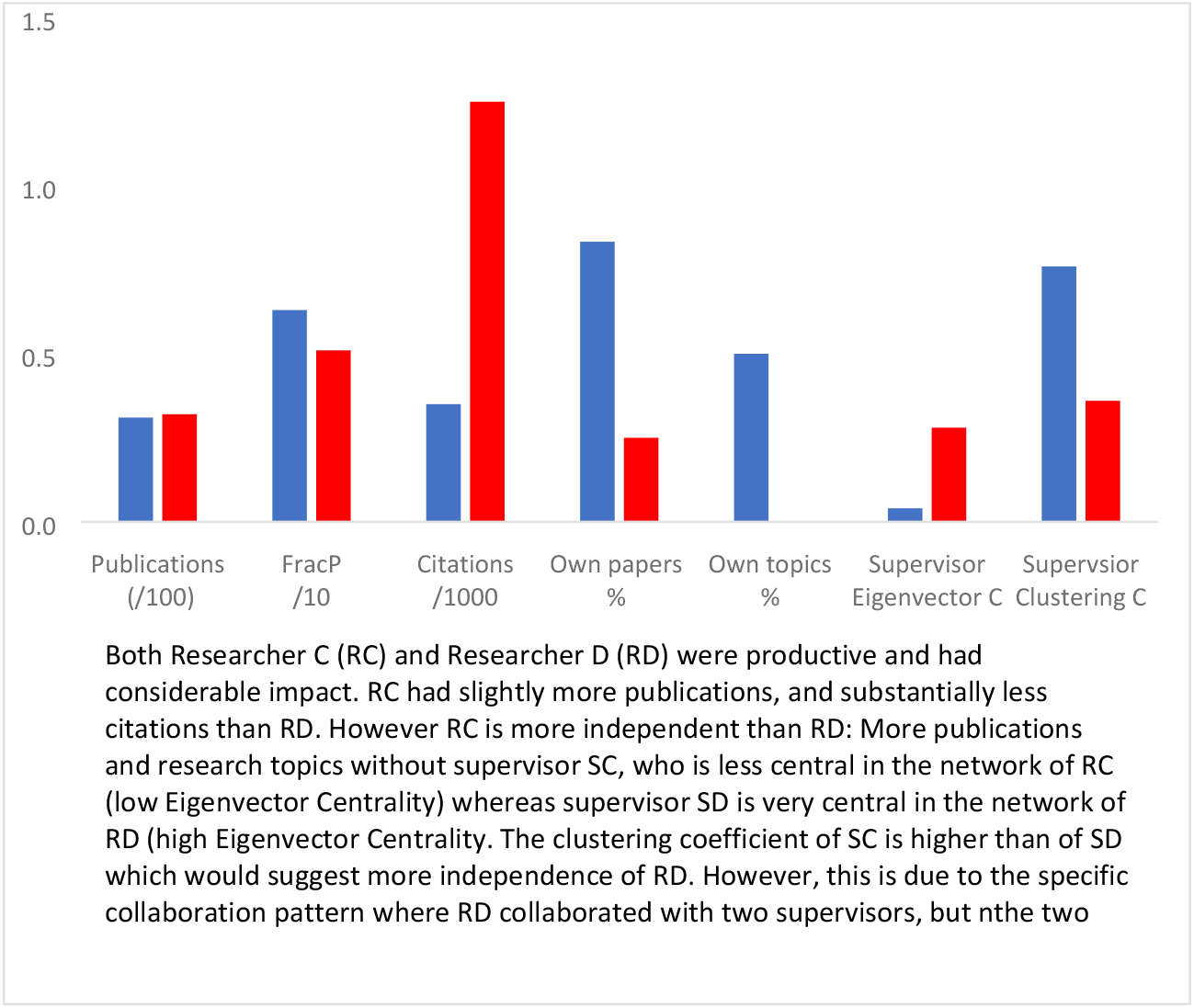
Indicator values for RC (blue) and RD (red)

It is interesting to have a deeper look at RD: despite having 25 % of the papers not coauthored with the two supervisors, these papers are in fact all in one of the research topics in which also the supervisors are active in (T10 and T12). This confirms that we indeed need more sophisticated indicators for independence than “some papers without the former supervisor” as e.g., the ERC formulates for the starting grant.

Then again, RC is also interesting from another perspective. RD has two supervisors, who not co-authored any paper. At the same time, we see RD modestly authoring and coauthoring with supervisor SD1 in topic T12, and frequent authoring and co-authoring with supervisor SD2 in topic T10 (Fig 5). Given this pattern, an alternative interpretation is that RD is not dependent on the supervisors, but doing original (interdisciplinary) work, linking the research lines of the two supervisors. Which of the interpretations is correct, is dependent on the moment of publishing: were RD’s papers in the two clusters published earlier than those of the supervisors or not? Inspecting the data suggests dependency, as the supervisors started to publish in those clusters years before RD entered those topics.

## Conclusions and discussion

This paper started by defining quality as a more-dimensional construct, and argued that there is a need for extending the indicators portfolio reflecting this multi-dimensionality. Then we argued that independence is an important dimension, which is emphasized in a variety of studies and reports. We proposed two indicators for this: the position of the former supervisor in the co-author network of a researcher, and the position of the former supervisor in the topic network of the researcher. These indicators could serve as a relevant start, as we have illustrated in the paper. The indicators also give a rather consistent picture: each of the different measures does put the researchers in about the same ranking, and the indicators together may constitute a valid measurement of independence. The overall (average) score clearly distinguish RA and RC from RB and RD (Table 4).

**Table 4:** Independence indicator scores.

| Rank order | Researcher | Overall score<sup>10</sup> | Centrality Supervisor (i) | Clustering Supervisor (ii) | Own papers (iii) | Own topics (iv) |
| --- | --- | --- | --- | --- | --- | --- |
| 1. Most independent | RC | 0.86 | 0.04 | 0.76 | 0.83 | 0.43 |
| 2. | RA* | 0.79 | 0.09 | 0.43 | 0.83 | 0.50 |
| 3. | RD** | 0.33 | 0.28 | 0.36 | 0.25 | 0.00 |
| 4. Least independent | RB | 0.13 | 0.91 | 0.11 | 0.04 | 0.14 |

\* We use here the first supervisor for calculating the indicators, the second supervisor gives similar values.
\*\* We use the second supervisor, as this supervisor was the main collaborator of RD (Fig 5).

Interestingly, RA had an equal output and a lower impact than RB, and RC had a somewhat lower output and a much lower impact than RD. But RA and RC had the successful academic career whereas the others stopped, indicating that the commonly discussed publication-based and citation-based indicators are not really informative in these career stages. At his career level, one assumes rather good numbers of publications and citations, and differences between those may not decisive in these career stages, whereas other evaluation dimensions than productivity and impact play a role. In our cases, the career success does correspond with the independence scores. We do not suggest that independence is always the decisive variable, as job selection is a multi-criteria decision-making problem.

In this era of *team science*, it is not anymore possible to show independence by single authored papers, therefore there is a need for more sophisticated assessment methods, and our proposal in this paper seems to be a step in that direction. Further development is needed, and several issues need to be addressed in order to have a functioning indicator:

i. Here we had four researchers in the chemistry and life sciences; and then using the WoS data seems a reasonable approach. However, as is often argued, for other fields the WoS is not covering enough of the output. Partly this is a statistical issue: If the WoS papers are representative for the whole oeuvre, also in terms of co-authorships, then relying on the WoS data may not be problematic. Nevertheless, we would agree that the indicators should be tested on other datasets too. For a social science case and a computer science case, we experimented with Google Scholar data, and this suggests that the proposed independence indicator can be generalized to other data sources [35].
ii. Another extension is to introduce a temporal dimension – one could imagine that an early career researcher starts a new topic without the supervisor(s) being involved, with the supervisors later jumping on that bandwagon when it proves to be successful. Then a joint topic is not a proof of dependence – on the contrary. This phenomenon was briefly discussed for researcher D and should be included in the indicators.
iii. Similarly, dependency may not only occur with the previous supervisor, but also with other collaborators later in the career. To control for that, the indicators should probably also be calculated for all frequent co-authors of the researcher.
iv. For individual evaluations, the data cleaning and calculations can be easily done. However, it would be useful to test on a large sample whether the independence indicator predict success better than other variables. In such a case data collection and cleaning may be resource and time intensive. That type of procedure indeed should be done for all indicators: their real life value should be shown in a multivariate prediction of career or grant success. If the independence indicators as defined here have validity, such test would show whether independence does make a difference.

Overall, this paper suggests that it is useful to derive more indicators for different quality dimensions of scholarly work. We would emphasize that this is a better way to move forward: if current indicators are not adequate (enough) as claimed e.g., in the Leiden Manifesto [8], one should try to develop better indicators, and not reduce the role of indicators and reinforce the role of peer review. Peer review is more problematic than indicators – as we argued in the introduction.

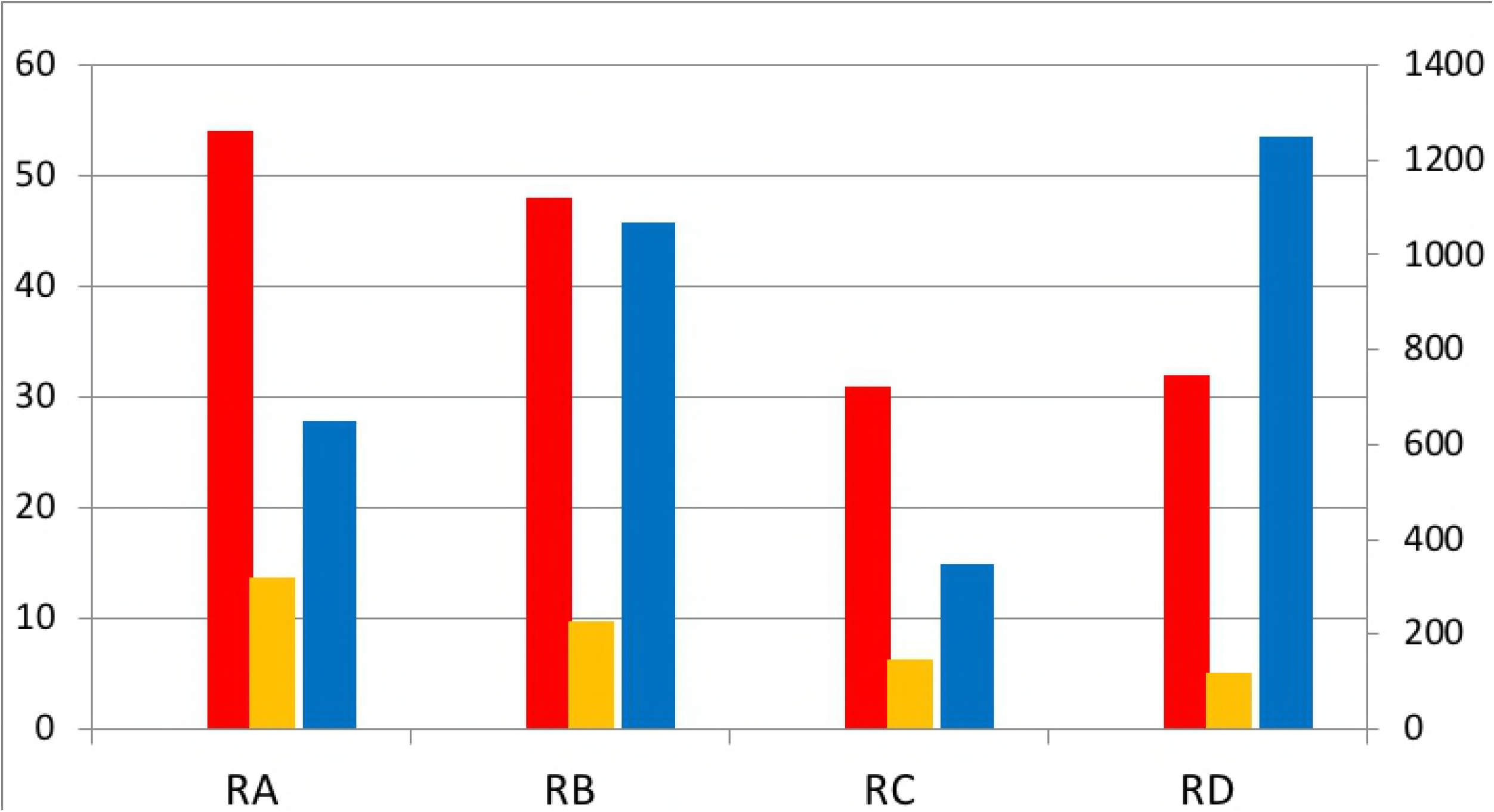

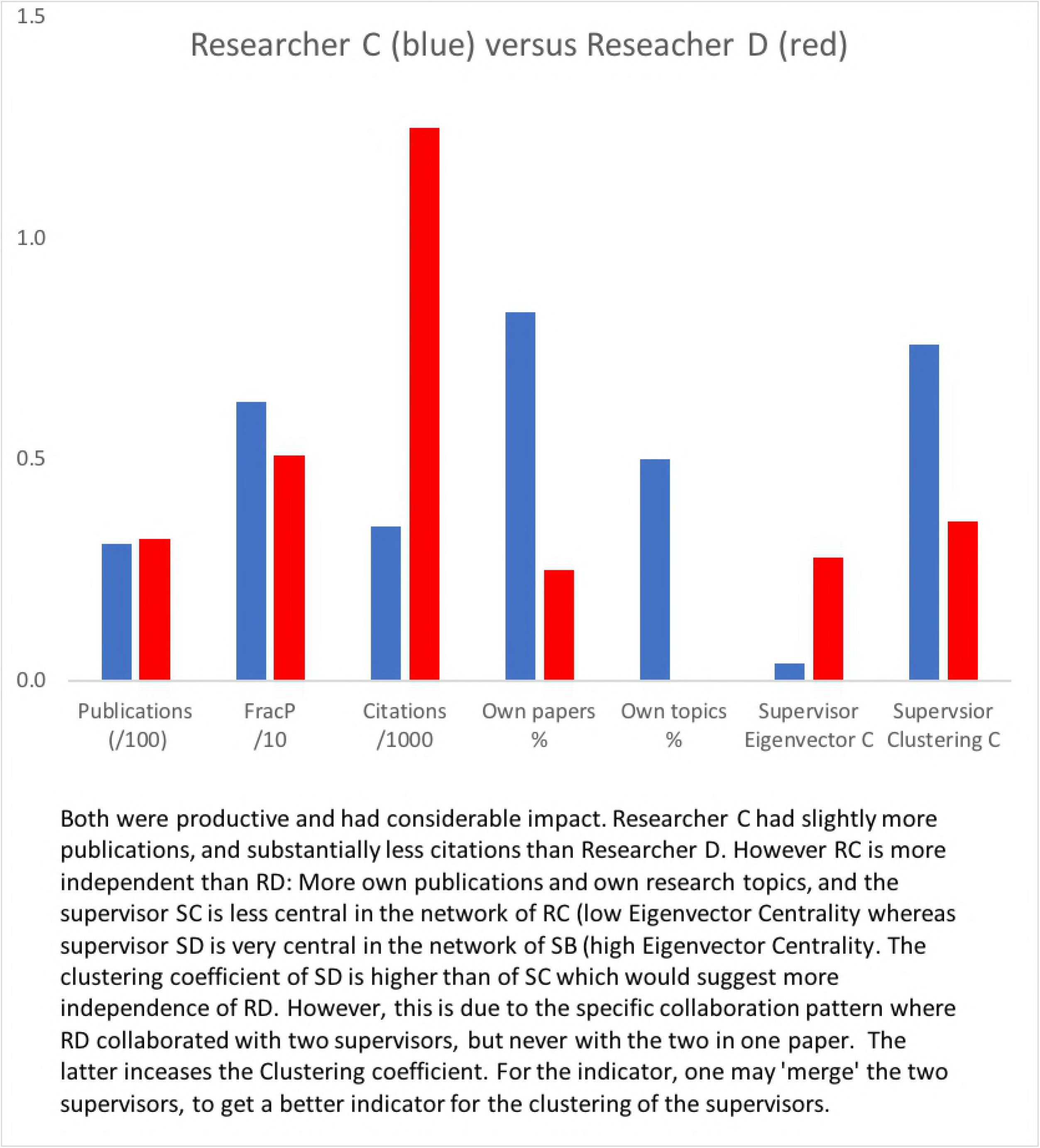

## Acknowledgements

An earlier version was presented at the STI conference 2012 in Montreal, Canada [36].

## Footnotes

1 Importantly, the critique on bibliometric indicators neglects the problems with peer review.

2 An unintended behavioral effect often mentioned is that researchers may start maximizing their indicator scores, instead of optimize their academic quality and impact. Yet, the often-claimed perverse effect of indicators (counting publications for example) lacks empirical support: Stimulating productivity is not detrimental to quality of research as Linda Butler claimed in her well known study on Australia (2003), but has a positive effect [37, 38].

3 Veira et al. indicates that bibliometric indicators do well as predictors of peer review in grant decisions [39]. However, in our view there is an obvious problem concerning mid-career researchers that are in the process of creating an independent career.

4 The idea is clear, but this indicator is easy to play: If supervisors know this, they will simply not be co-author anymore on all papers. Indeed, in CV analysis we see that the applicants lists papers without the supervisor, but further inspection shows that some are with the co-supervisor. Another requirement is the ability to do groundbreaking research. This can be measured by (i) having published top cited papers, and (ii) doing research within promising and hot topics, indicated by e.g., fast growth.

5 Other indicators mentioned are: being PI on a grant; being lab director or research leader; evaluations of teaching for being sole teacher; letters from research directors about your contribution to a research project (including assigning you a percentage of effort)

6 The Swedish rank ‘docent’ is equal to the US rank “associate professor” or the British “research fellow”, but is a non-paid title indicating your level of academic competence in a certain field.

7 The STEM fields are mainly producing co-authored papers. In the social sciences and humanities this is increasing, but here single authored papers are rather frequent. For those fields, we may have to develop a different type of independence indicator.

8 In a limited number of topics, the researcher and former supervisor did not co-author, but the supervisor was marginally active a decade before the researcher entered these topics. We do not consider this as a shared topic.

9 As most of the early career of the researchers under study took part in the 1990s, we do not have the data for calculating field normalized (size dependent and size independent) indicators such as visibility in the top 10% cited papers. However, as the researchers work in similar fields, this is for the current analysis not a problem.

10 Calculated as [(1 − i) + ii + iii + 2 * iv] / 4. The score of (iv) is weighted double, as the range is between 0 and 0.5 whereas the other indicates have a range between 0 and 1.

